# FOXP4 differentially controls cold-induced beige adipocyte differentiation and thermogenesis

**DOI:** 10.1101/2021.10.11.463898

**Authors:** Fuhua Wang, Shuqin Xu, Tienan Chen, Shifeng Ling, Wei Zhang, Shaojiao Wang, Rujiang Zhou, Xuechun Xia, Zhengju Yao, Pengxiao Li, Xiaodong Zhao, Jiqiu Wang, Xizhi Guo

## Abstract

Beige adipocytes possess a discrete developmental origin and notable plasticity in thermogenic capacity in response to various environmental cues. But the transcriptional machinery controlling beige adipocyte development and thermogenesis remains largely unknown. By analyzing beige adipocyte-specific knockout mice, we identified a transcription factor, Forkhead Box P4 (FOXP4) that differentially governs beige adipocyte differentiation and activation. Depletion of *Foxp4* caused a decline in the frequency of beige preadipocytes by switching their cell fate towards fibroblastic cells at the expense of beige adipocytes. However, we observed that ablation of *Foxp4* in differentiated adipocytes profoundly potentiated their thermogenesis upon cold exposure. Of note, the outcome of *Foxp4*-deficiency on UCP1-mediated thermogenesis was confined to beige adipocytes, rather than to brown adipocytes. Taken together, we submit that FOXP4 primes beige adipocyte cell fate commitment and differentiation by potent transcriptional repression of the thermogenic program.

## INTRODUCTION

Beige adipocytes, a population of thermogenic adipocytes distinct from brown adipocytes, have recently caught mainstream attention. Their relevance to adult humans holds promise as a new therapeutic target in combating obesity and other metabolic disorders. Beige adipocytes emerge within white adipose tissue (WAT) depots in response to various environmental cues, including chronic cold acclimation, exercise, β3-adrenergic receptor (AR) agonists, cancer cachexia, and tissue injury (Barbatelli et al., 2010; Rosenwald et al., 2013). As with classical brown adipocytes, beige adipocytes present with multilocular morphology and produce heat mainly through UCP1-mediated thermogenesis. Lineage tracing studies demonstrated that beige adipocytes consist of heterogeneous subpopulations of distinct origins depending on the nature of external induction stimuli (Berry et al., 2016). For example, beige adipocytes originate either from PDGFRα^+^ stromal progenitor cells (Han et al., 2021; Lee et al., 2015; Lee et al., 2012), or from mural perivascular cells when targeted by *SMA-CreERT* (Long et al., 2014). These two beige progenitor populations are not mutually exclusive. A recent study noted that PDGFRα^+^ progenitor cells are required for developmental adipogenesis, but not for adult beige adipogenesis (Shin et al., 2020). Further evidence showed that beige precursor cells are a population distinct from brown or white fat precursors with selective cell surface markers, including CD81 (Oguri et al., 2020) and CD137 (Wu et al., 2012).

Following cell fate determination and differentiation, beige adipocytes possess inducible and reversible thermogenic capacity, as well as plasticity in cellular morphology in response to environmental stimuli (Paulo and Wang, 2019). The first wave of *de novo* beige adipocytes express UCP1 with multilocular morphology. They appear in WAT of mice independent of temperature conditions at the peri-weaning stage of development (Wu et al., 2020). At the adult stage, these beige adipocytes regress, lose UCP1 expression and are morphologically identical to white adipocytes at room temperature. The dormant beige adipocytes can reappear in response to cold challenge, and reverse back to white-like cellular morphology at conditions of re-warming or withdrawal of the β3-AR agonist (Roh et al., 2018; Rosenwald et al., 2013). Previous studies have proposed that cold-induced beige adipocytes predominantly arise from transdifferentiation of preexisting white adipocytes (Barbatelli et al., 2010; Cattaneo et al., 2020; Rosenwald et al., 2013). However, this view requires cautious assessment because of the current technical limitations to determine how many of those pre-existing “white” adipocytes are *de facto* latent beige adipocytes with UCP1^+^ history. Of note, the conversion between thermogenically latent and active states in beige fat cells may be attributed to reversible processes of mitochondrial biogenesis and clearance (Altshuler-Keylin et al., 2016), as well as to chromatin reprogramming and function of specific transcriptional machinery (Roh et al., 2018).

Despite these differences in developmental origins and physiological functions of brown, beige and white adipocytes, these cell types share a similar transcriptional cascade that controls the process of fat cell differentiation. Several factors are commonly employed by both brown and beige adipocytes, including Cebpα/β-PPARγ cascades for early cell fate commitment and differentiation, as well as activators of EBF2, Prdm16 and PGC1α for thermogenesis (Shapira and Seale, 2019; Wang and Seale, 2016). However, as aforementioned, beige adipocyte plasticity in cell differentiation and thermogenesis involves a cell-specific changes in morphology, transcription, and chromatin landscape. In contrast to our significant understanding of brown adipocyte transcription, the beige-selective regulatory machinery remains largely unclear.

Forkhead Box P4 (FOXP4) typically functions as a transcription factor that regulates islet α cell proliferation (Spaeth et al., 2015), breast cancer invasion (Ma and Zhang, 2019), and speech/language (Snijders Blok et al., 2021). Our previous studies have shown that FOXP4 also controls endochondral ossification via a complex with FOXP1 and FOXP2 (Zhao et al., 2015). In this study, we employed *SMA-Cre*^*ERT*^ and *AdipoQ-Cre* mice to knockout *Foxp4* in beige precursors and differentiated beige cells, respectively. Inactivation of *Foxp4* in progenitor cells impaired beige fat cell differentiation and switched them to pro-fibrotic cell potency. In contrast, *Foxp4* deficiency in differentiated beige adipocytes exacerbated their cold-induced thermogenesis. Mechanistically, we found that FOXP4 directly repressed transcription of *PDGFRα, PGC1α* and *Cebpβ*, thereby acting as a transcriptional “brake” on these critical components of beige adipocyte regulation. Together, our results suggest that FOXP4 not only primes early cell fate commitment of beige adipocytes, but also attenuates their cold-induced thermogenesis.

## RESULTS

### Dynamical expression of *Foxp4* during beige adipocyte differentiation

To investigate the expression pattern of FOXP4 in adipose tissues, two representative subpopulations of adipocytes, interscapular brown adipose tissues (BAT) and subcutaneous white adipose tissues (sWAT), were obtained from 8-week-old wild-type C57BL/6J mice that were housed at room temperature (23 °C) . High levels of FOXP4 expression were detected within brown and white adipocytes by immunofluorescence analyses (Fig. 1A). Next, stromal vascular fraction (SVF) cells isolated from sWAT of wild type mice were induced to beige adipocytes *in vitro* by culturing in beige adipogenic media for 7 days. As expected, *PPARγ* and *Ucp1* expression levels were elevated during beige adipocyte differentiation, whereas *Foxp4* expression peaked at day 2 of induction and declined swiftly thereafter (Fig. 1B). In addition, western blotting and qPCR analysis revealed that FOXP4 expression in BAT and sWAT was slightly increased in response to cold exposure (Supplementary Fig. S1A, B). These dynamic alterations in expression implicated a phase-specific function of FOXP4 during beige adipocyte differentiation.

**Fig. 1.**
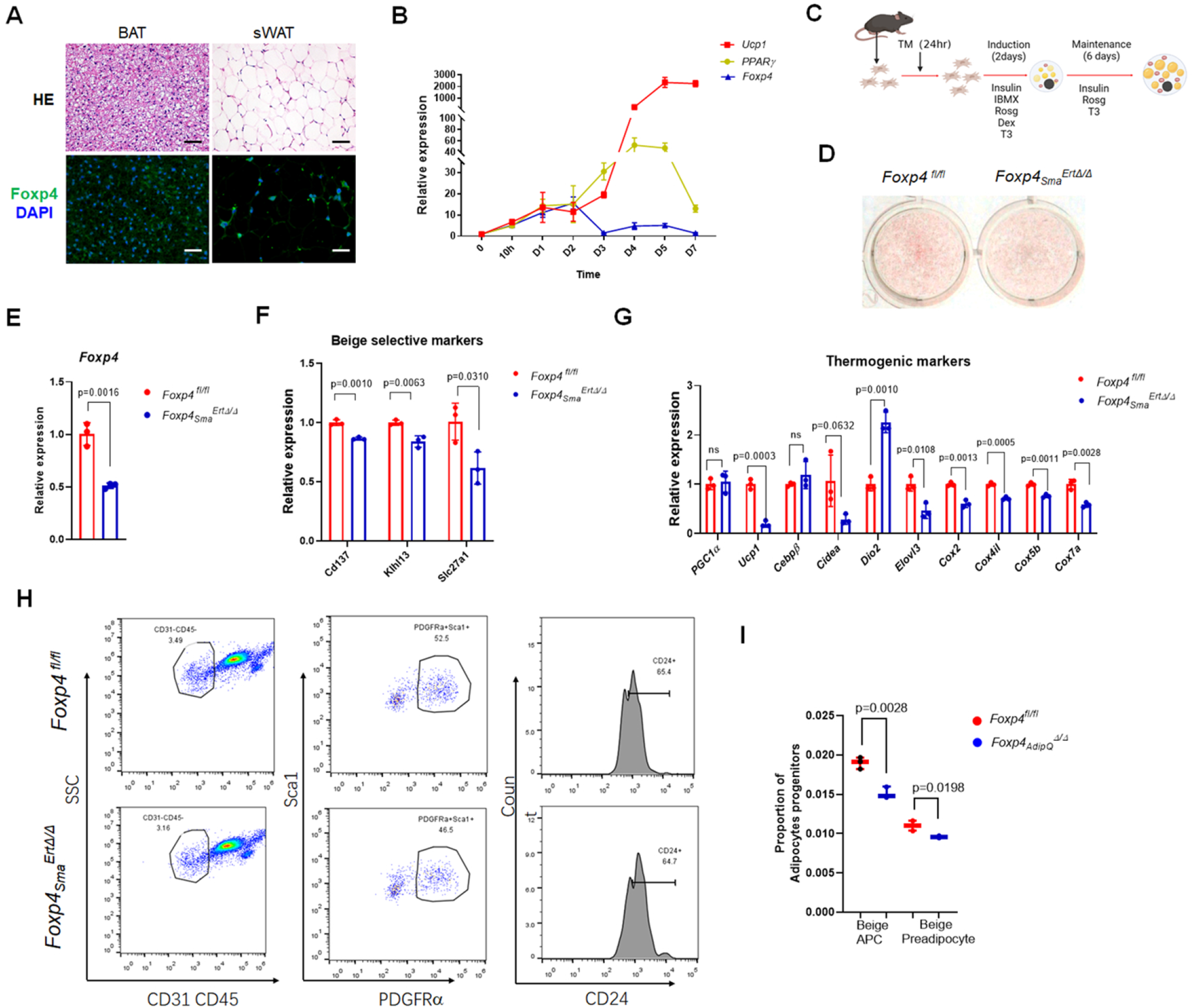
Ablation of *Foxp4* impairs beige adipocyte differentiation. (A) Hematoxylin and eosin (H&E) staining and immunofluorescence examination for FOXP4 on sections from BAT and sWAT of 2-month-old wild type mice. Bar, 100μm. (B) mRNA levels of *Foxp4, PPARγ, Ucp1* during culture courses of beige adipocyte differentiation derived from stromal vascular fraction (SVF) of sWAT depot. n, 3. (C) Diagram depicting that sWAT-SVF cells were treated with tamoxifen (TM) 24-hour post isolation and cultures to induce *Foxp4* knockout in beige cells, which were designated as *Foxp4*^*fl/fl*^ and *Foxp4*_*Sma*_^*ErtΔ/Δ*^. Then these precursor cells underwent 8-day beige induction cultures. (D) Oil Red O staining for SVF-derived beige adipocytes from (C). (E-G) mRNA levels of *Foxp4*, beige selective and thermogenic markers in cells from (C). n, 3. (H) Flow cytograms showing expression of for beige adipocyte progenitor cells (APC, CD31^-^CD45^-^PDGFRα^+^Sca1^+^CD24^-^) and beige preadipocytes (CD31^-^CD45^-^ PDGFRα^+^Sca1^+^CD24^+^) 24-hour post tamoxifen induction in SVF cell cultures. (I) Quantitative analysis for proportion of beige adipocyte progenitor cells and beige preadipocytesin (H). n, 3.

### Ablation of *Foxp4* impairs beige adipocyte differentiation

A major proportion of adult beige adipocytes is reported to be derived from mural progenitor cells within sWAT, which could be selectively and conditionally targeted by *SMA-Cre*^*ERT*^ (Long et al., 2014). Tamoxifen-inducible *Foxp4* conditional knockout mice (thereafter designated as *Foxp4*_*Sma*_^*ErtΔ/Δ*^) were generated by crossing *Foxp4*^*fl/fl*^ with *SMA-Cre*^*ERT*^ mice. SVF cells obtained from sWAT of *Foxp4*_*Sma*_^*ErtΔ/Δ*^ were maintained for 2 days within cultures with tamoxifen, followed by another 6-days in beige adipogenic differentiation cultures (Fig. 1C). As shown in Fig. 1E, beige cell differentiation was impaired following loss of *Foxp4*, as evidenced by oil red staining (Fig. 1D). In addition, we observed decreased expression levels of *Foxp4* (Fig. 1E), beige-specific (*CD137, Klh13, Slc27a1*) (Fig. 1F), and thermogenic marker (*PGC-1α, Ucp1, Cebpβ, Cidea, Dio2, Elovl3, Cox2, Cox4il, Cox5b, Cox7a*) genes (Fig. 1G).

The defects of beige cell differentiation may stem from perturbed *de novo* biogenesis. We next tracked CD29^+^ PDGFRα^+^ Sca1^+^CD24^+^ adipocyte progenitor cells (APC) within WAT-SVF 24-hour post tamoxifen administration by flow cytometry (FACS) as described previously (Berry and Rodeheffer, 2013; Lee et al., 2015). The frequency of beige APC (defined as CD31^-^CD45^-^PDGFRα^+^Sca1^+^) and beige preadipocyte (defined as CD31^-^CD45^-^PDGFRα^+^Sca1^+^CD24^+^) were relatively lower in SVF cells o*Foxp4*_*Sma*_^*ErtΔ/Δ*^ as compared to that of *Foxp4*^*fl/fl*^ (Fig. 1H, I). These observations indicated that FOXP4 inactivation impaired beige adipocyte differentiation, potentially the early beige cell commitment.

### Deletion of *Foxp4* switches stromal progenitor cells towards fibroblasts at the expense of beige adipocytes

FACS analysis indicates the potential role of FOXP4 in beige cell early commitment. PDGFRα^+^ stromal progenitor cells also give rise to beige adipocytes during early development (Gao et al., 2018), also have the potential to adopt fibroblastic cell fate upon activation of the PDGFRα pathway (Shin et al., 2020; Sun et al., 2017). Unfortunately, when *Foxp4* was inactivated by *SMA-Cre*, the knockout mice showed defects in postnatal development and growth (data not shown). This blocked our ability to investigate FOXP4 function during beige cell development. To circumvent this problem, we transfected the SVF cells from *Foxp4*^*fl/fl*^ sWAT with either retroviral *pMSCV-Cre* or, as a control, retroviral *pMSCV-GFP*. This approach allowed us to achieve efficient and extensive inactivation of *Foxp4* in progenitors. As shown in Fig. 2A-C, beige adipocyte differentiation in *Foxp4*-deficient stromal progenitor cells (designated as *Foxp4*_*pMSCV*_^Δ/Δ^) was impaired, as evidenced by oil red staining and down-regulation of a set of thermogenic (*PPARγ, PGC-1α, Adrb3, Ucp1, Cidea, Dio2, Elovl3*) and beige-elective signature genes (*CD137, Tbx1, Tmem16, Slc27a1*). Similar, but less penetrant alterations in thermogenesis was were detected in BAT-SVF cells (Supplementary Fig. S2A-C). In contrast, deletion of *Foxp4* by *pMSCV-Cre* had little effect on general adipogenesis, as evidenced by Oil red staining and qPCR analysis based on sWAT-SVF adipogenic cultures without T3 and TZDs (Supplementary Fig. S2E, F).

**Fig. 2.**
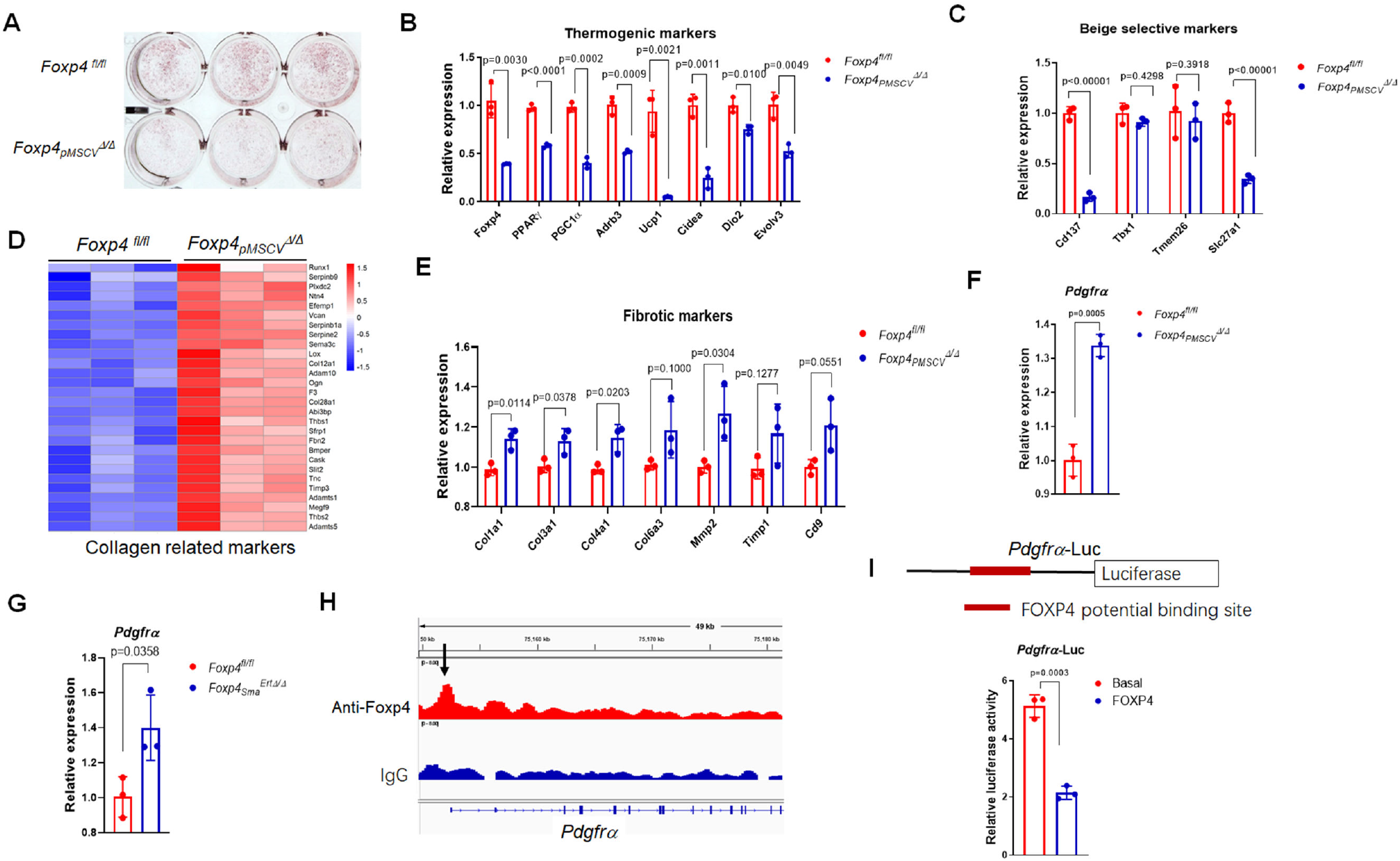
*Foxp4* deficiency disrupts beige-fibroblast balance in progenitor cells through regulating *Pdgfrα* expression. A) Primary precursor cells from sWAT of *Foxp4*^*fl/fl*^ mice were transfected with retrovirus of *pMSCV-Cre* or *pMSCV-GFP* to induce *Foxp4* inactivation before being induced to undergo beige adipocyte differentiation. Oil Red O staining was performed 8 days post differentiation cultures. B, C) mRNA levels of thermogenic and beige selective markers in cells of (A). n, 3. D) Heatmap depicting the mRNA levels of collagen related markers in beige adipocytes in (A). E) mRNA levels of fibrotic markers in beige cells of (A). n, 3. F, G) qPCR validated the increased *Pdgfrα* expression levels in beige adipocytes with *Foxp4* deficiency by *pMSCV-Cre* (*F*) or *SMA-CreER* (G). n, 3. H) Chromatin occupancy analysis by ChIP-seq showed the relative enrichment of Foxp4 binding sites (black arrows) upstream of *Pdgfrα* gene promoter region, based on beige adipocytes derived from SVF of wild type sWAT. I) Luciferase reporter assay validated the repressive activity of Foxp4 protein in *Pdgfrα* gene transcription. n, 3. The upper panel depicts the *Pdgfrα*-Luc construct and potential FOXP4 binding site.

Interestingly, RNA-seq analysis revealed that an array of collagen fibril-related transcripts were relatively enriched in *Foxp4*_*pMSCV*_^Δ/Δ^ beige cells as compared to controls (Fig. 2D). Elevated expression levels of several of these pro-fibrotic genes (*Col1a1, Col3a1, Col4a1, Col6a3, Mmp2, Timp1, CD9*) were validated by qPCR analysis (Fig. 2E). Accordingly, similar alterations of profibrotic marker genes were observed in beige cells obtained from sWAT-SVF of *Foxp4*_*Sma*_^*ErtΔ/Δ*^ mice following *Foxp4* inactivation induced by tamoxifen (Supplementary Fig. S2D). Thus, FOXP4 expression is required to balance fibroblast-beige ratios in stromal progenitor cells.

Given the unexpected finding that high levels of PDGFRα in stromal progenitor cells switch beige adipocytes to fibroblasts, we then investigated the impact of FOXP4 on *Pdgfrα* gene transcription. As shown in Fig. 2F and G, qPCR analysis revealed that *Pdgfrα* transcripts were increased in *Foxp4*_*pMSCV*_^Δ/Δ^ or in *Foxp4*_*Sma*_^*ErtΔ/Δ*^ cells. In addition, Chip-seq analysis of SVF progenitor cells detected relatively high enrichment of FOXP4 binding sites within the promoter region of *Pdgfrα* (arrow in Fig. 2H). Luciferase reporter assays employing a *Pdgfrα* promoter-driven luciferase as substrate validated that FOXP4 repressed transcription of *Pdgfrα* in 293T cells (Fig. 2I). Together these data suggested that FOXP4 controls beige adipocyte cell fate commitment from progenitors by directly modulating *Pdgfrα* transcription.

### Ablation of *Foxp4* modestly augments juvenile and mature beige adipocyte thermogenesis

The first wave of *de novo* beige adipocyte biogenesis and UCP1 activation occurs within sWAT at peri-weaning stage independent of temperature conditions (Wang et al., 2017; Wu et al., 2020). To evaluate the potential impact of *Foxp4* deficiency on beige adipocyte thermogenesis, we eliminated *Foxp4* in differentiated adipocytes with *AdipoQ-Cre* (Eguchi et al., 2011), hereafter designated as *Foxp4*_*AdipQ*_^Δ/Δ^. We confirmed (Supplementary Fig.S3A, B) that FOXP4 was efficiently reduced at the mRNA and protein levels in BAT and sWAT from *Foxp4*_*AdipQ*_^Δ/Δ^ mice. *Foxp4*_*AdipQ*_^Δ/Δ^ mice appeared relatively normal in size and in fat depots of BAT and sWAT at 3 weeks of age (Fig. 3A, B). H&E staining and IHC analysis revealed that higher numbers of UCP1^+^ beige adipocytes resided in sWAT from *Foxp4*_*AdipQ*_^Δ/Δ^ knockout mice than in controls (Fig. 3C). Consistent with that observation, qPCR analysis confirmed the up-regulation of a set of thermogenic and mitochondrial signature genes (*Cebpβ, Cidea, Dio2, Elovl3, PGC-1α, Ucp1, Cox2, Cox4il, Cox5b, Cox8b*) (Fig. 3D, E). However, the expression of several beige selective marker genes (*CD137, CD40, Klh13, Tbx1*) showed no obvious increases (Fig. 3F). These observations suggested that FOXP4 boosted juvenile beige adipocyte activation, but had little effect on their early differentiation.

**Fig. 3.**
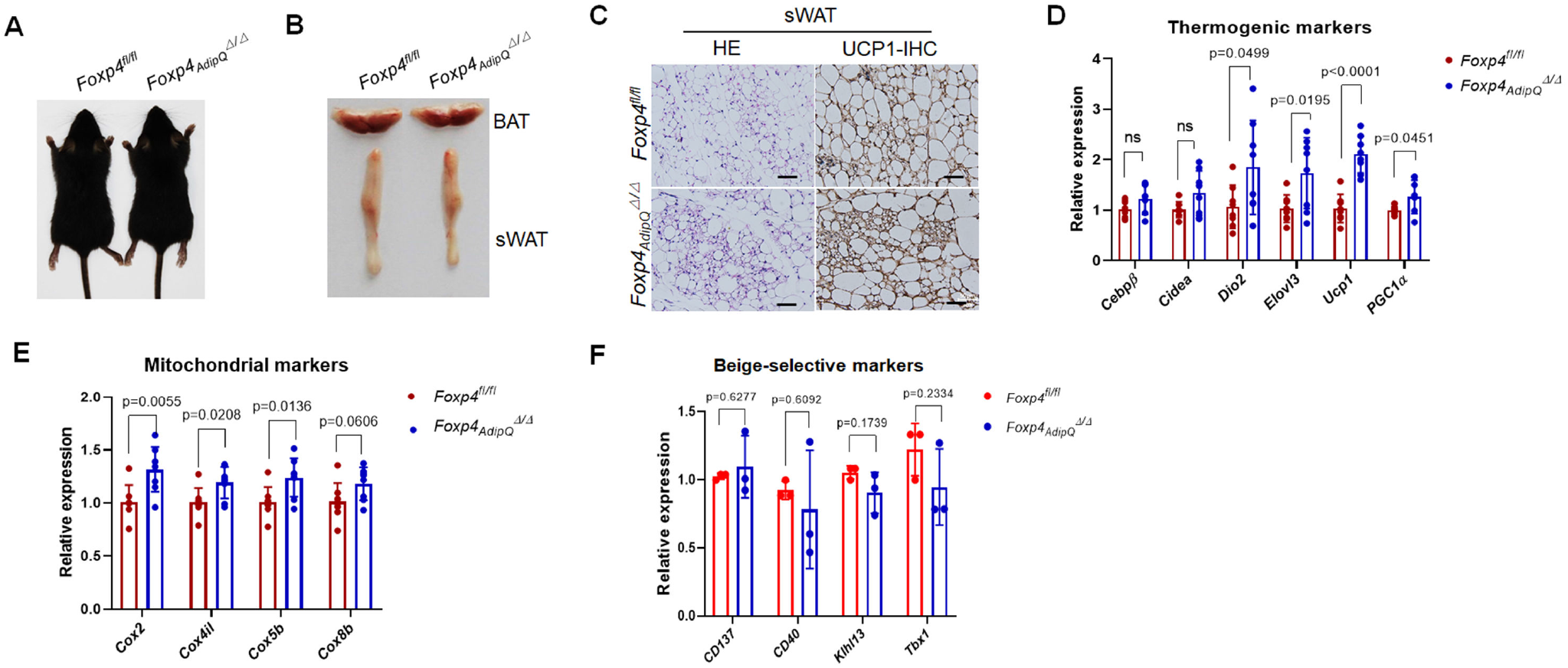
*Foxp4* deletion mildly augments beige adipocytes thermogenesis at peri-weaning stage. A) Dorsal view of representative *Foxp4*^*fl/fl*^ and *Foxp4*_*AdipQ*_ ^Δ/Δ^ mice at 3 weeks old. B) Gross morphology of BAT and sWAT depot of mice (A). C) H&E and immunohistochemical staining (IHC) for UCP1 on sWAT sections from *Foxp4* _*AdipQ*_^Δ/Δ^ mice of (*A*). D, E) mRNA levels of thermogenic and mitochondrial markers in sWAT. F) qPCR validated the relative normal expressions of beige selective marker genes in sWAT from *Foxp4* _*AdipQ*_^Δ/Δ^ mice.

The transcriptional pathway underlying adult beige cell differentiation and thermogenic activation is distinct from that of the juvenile at the peri-weaning stage (Wu et al., 2020). *Foxp4*_*AdipQ*_ ^Δ/Δ^ knockout mice appeared normal in size, weight and adiposity compared to controls at adult stages (Fig. 4A). They were modestly smaller at 5 months (Fig. 4B). Accordantly, the thermogenic program in the BAT of *Foxp4*_*AdipQ*_ ^Δ/Δ^ mice did not appear to be activated at ambient temperature (20-22 °C), as indicated by H&E staining and IHC, as well as by qPCR analysis of a set of thermogenic marker transcripts (Supplementary Fig. S3C, D). Indirect calorimetry analysis by CLAMS revealed that *Foxp4* loss had no impact on O_2_ or CO_2_ consumption, as well as on energy expenditure following identical diet and locomotor activity as controls at 20 ∼22 °C (Supplementary Fig. S4). In contrast, thermogenic activation of beige adipocytes was mildly exacerbated in sWAT of *Foxp4* _*AdipQ*_^Δ/Δ^ mice, as evidenced by relatively enriched levels of UCP1^+^ adipocytes and by upregulation of thermogenic or mitochondrial signature genes (*PPARγ, Dio2, Cidea, Ucp1, Cox2, Cox4il, Cox5b, Cox8b*) (Fig. 4C, D). We conclude from these analyses that loss of FOXP4 results in modest augmentation of thermogenesis in beige adipocytes at ambient temperature.

**Fig. 4.**
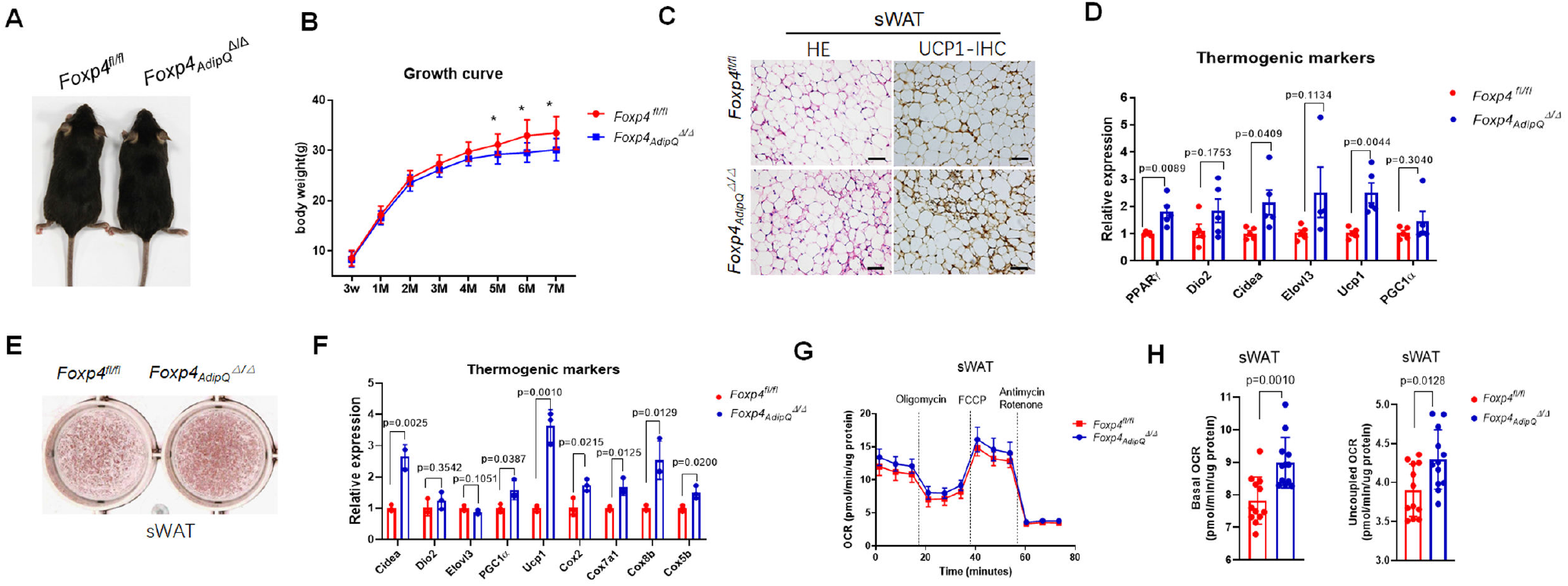
FOXP4 cell-autonomously repressed beige adipocytes thermogenesis. A) Representative dorsal view *Foxp4*^*fl/fl*^ and *Foxp4*_*AdipQ*_^Δ/Δ^ mice at age of 2 months old. B) Growth curve showed that the body weights were mildly decreased in *Foxp4*_*AdipQ*_^Δ/Δ^ mice as compared to that of *Foxp4*^*fl/fl*^ control mice since age of 5 months old. n, 7. (C) H&E and IHC staining for UCP1 protein on sWAT sections from *Foxp4*_*AdipQ*_^Δ/Δ^ mice at age of 2 months old. Bar, 100μm. (D) mRNA levels of a set of thermogenic genes in sWAT of *Foxp4*_*AdipQ*_^Δ/Δ^ mice. n, 3. E) Oil Red O staining 8 day post brown adipocyte differentiation from sWAT-SVF of *Foxp4*^*fl/fl*^ and *Foxp4*_*AdipQ*_^Δ/Δ^ mice at age of 8 weeks. F) mRNA levels of thermogenic markers for beige adipocytes in (A). n, 3. G) Oxygen consumption rate (OCR) was measured for beige adipocytes from (E). Uncoupled respiration was recorded after oligomycin inhibition of ATP synthesis, and maximal respiration following stimulation with carbonyl cyanide 4-(trifluoromethoxy) phenylhydrazone (FCCP). n, 3. H) Quantitative analysis of basal and uncoupled OCR in (G). n, 3.

### *Foxp4* deficiency protects mice from HFD-induced obesity

Beige adipocyte thermogenesis could combat obesity in human (Lidell et al., 2013) . To examine the impact of *Foxp4* deficiency on long-term adipose tissue metabolism, mice were fed with HFD for three months. We observed that *Foxp4*_*AdipQ*_^Δ/Δ^ mice were leaner in body, with fewer adipose depots (Supplementary Fig. S4A, B), and gained less body weight and adiposity than littermates after 12-week HFD feeding starting at ages of 2 months (Supplementary Fig. S4C). *Foxp4*_*AdipQ*_^Δ/Δ^ mutant mice also retained more efficient glucose tolerance following HFD feeding, as evidenced by GTT scores (Supplementary Fig. S4D). Of note, as aforementioned, UCP1-mediated beige thermogenesis was mildly mitigated in *Foxp4*_*AdipQ*_^Δ/Δ^ mutant mice at room temperature (Fig. 3C, D and Fig. 4C, D), which may undermine the effect that *Foxp4*-deficiency protects from HFD-induced obesity.

### FOXP4 suppresses beige adipocyte thermogenesis cell-autonomously

To examine a potential cell-autonomous effect of FOXP4 on beige adipocyte energy metabolism, SVF cells were isolated from sWAT depots of *Foxp4* _*AdipQ*_^Δ/Δ^ and control mice and then induced for beige adipocyte differentiation *in vitro*. We observed advanced beige adipocyte thermogenesis in *Foxp4*-deficient SVF progenitors, as evaluated by elevated mRNA levels of a set of thermogenic genes (*Cidea, Dio2, Elovl3, PGC-1α, Ucp1, Cox2, Cox7a1, Cox5b, Cox8b*) (Fig. 4E,F). In addition, oxygen consumption rates (OCR) of beige adipocytes from *Foxp4*_*AdipQ*_ ^Δ/Δ^ mutant mice exhibited higher total and uncoupled OCR as compared to controls (Fig. 4G, H). These results indicated that loss of FOXP4 led to elevated mitochondrial respiration. The thermogenic potency of SVF-derived brown adipocyte also was potentiated in *Foxp4*-deficient mice, as evidenced by qPCR and OCR analyses (Supplementary Fig. S3E-H). This line of evidence indicated that FOXP4 suppresses beige adipocyte thermogenesis in a cell-autonomous manner.

### *Foxp4* depletion exacerbates cold-induced beige adipocyte thermogenesis *in viv*

We demonstrated that beige adipocytes could be mildly activated within sWAT under room temperature at adult stage. Then we examined beige adipocyte thermogenesis and activation in *Foxp4*_*AdipQ*_^Δ/Δ^ mice under cold conditions. As compared to controls, *Foxp4* knockout mice had relatively higher rectal temperature during 6-hour 4 °C exposure (Fig. 5A). *Foxp4*_*AdipQ*_^Δ/Δ^ mice also displayed a “browning” feature of sWAT after one-week at 4 °C (Fig. 5B). This result was consistent with H&E staining and IHC results demonstrating higher levels of UCP1^+^ beige adipocytes (Fig. 5C). Elevated beige adipocyte thermogenesis was validated by RT-qPCR analysis of signature genes (*Cidea, Dio2, Elovl3, PGC-1α, Ucp1, Cox2, Cox5b, Cox8b*) (Fig. 5D). Consistent with these observations, transmission electronic microscopic (TEM) and qPCR analyses revealed relative enrichment of mitochondria within beige adipocytes of *Foxp4*_*AdipQ*_^Δ/Δ^ sWAT (Fig. 5F, G). Yet long-term cold exposure had no significant effect on beige adipocyte *de novo* biogenesis in mutant mice, as evidenced by transcript levels of beige signature genes (*Klh13, Sl27a1, Tbx1, CD137*) (Fig. 5E). We further observed that cold exposure had little effect on thermogenesis in the BAT of *Foxp4* _*AdipQ*_^Δ/Δ^ mice (Supplementary Fig. S6). Together, these results suggested that FOXP4 acts as a repressor of the thermogenic gene program in beige adipocytes.

**Fig. 5.**
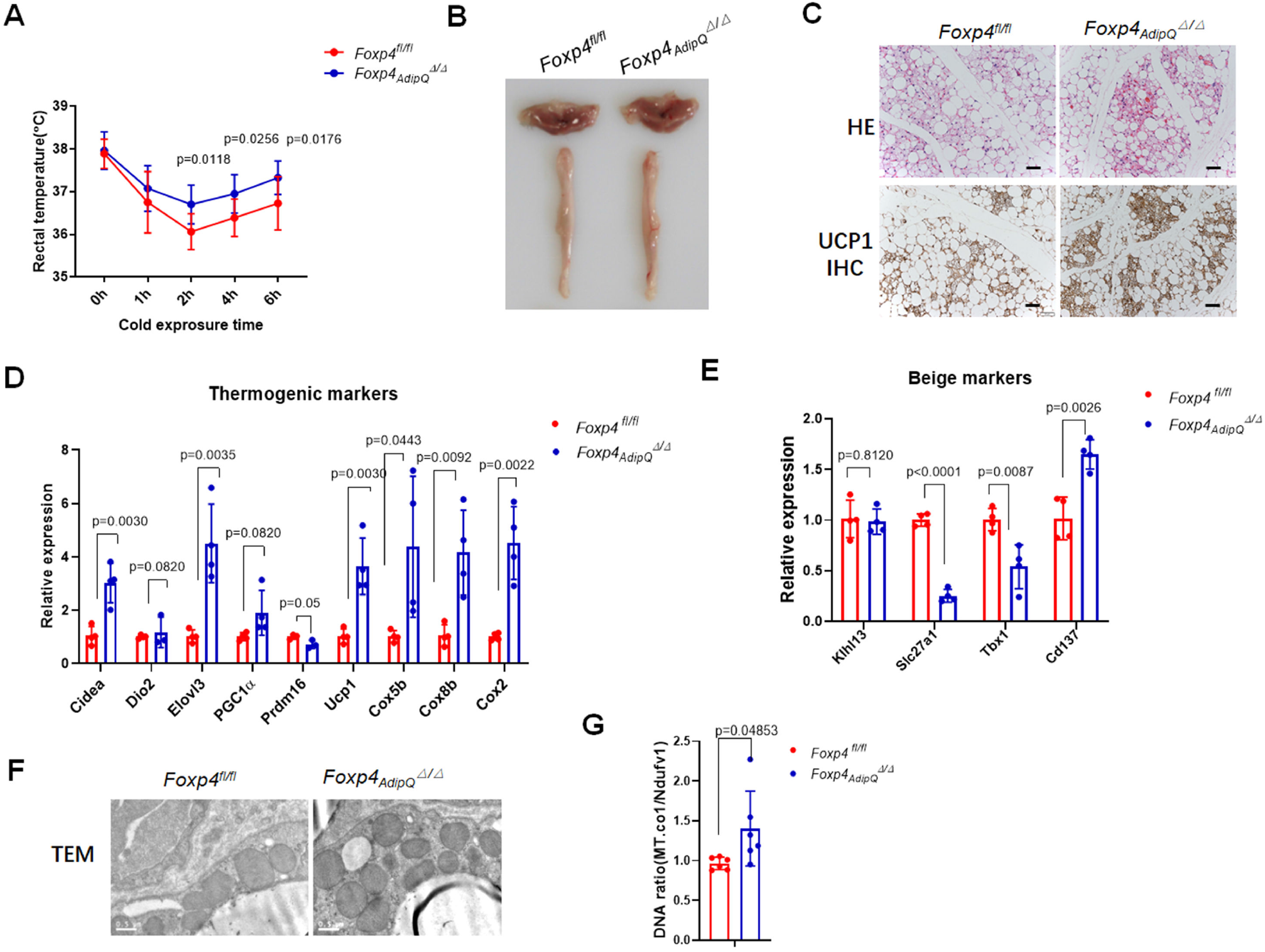
Loss of *Foxp4* exacerbated beige adipocytes thermogenesis upon cold exposure. A) Record of rectal temperature of 2-month-old *Foxp4*_*AdipQ*_ ^Δ/Δ^ mice during 6-hour 4 °C cold challenge. n, 8. B) Fat depot of BAT and sWAT in *Foxp4*_*AdipQ*_^Δ/Δ^ mice after one-week 4 °C exposure. C) H&E and IHC staining for UCP1 on sWAT sections from mice of (B). Bar, 100μm. D, E) mRNA levels of a set of thermogenic and beige selective genes. n, 3. F) Transmission electronic micrographs (TEM) of sWAT from mice (B). Bar, 2μm. G) Mitochondria DNA abundance in sWAT of mice (B). n, 4.

Thermogenesis of beige adipocytes could also be activated through adrenergic signaling (Jiang et al., 2017). However, when treated with 7-day consective injection of the β3-AR agonist, CL-316,243, *Foxp4*_*AdipQ*_^Δ/Δ^ mice exhibited no evident activation of thermogenic program nor no increase in beige adipocyte biogenesis, as determined by H&E staining, IHC and RT-qPCR analyses (Supplementary Fig. S7). These findings suggested that FOXP4 controls cold-induced beige adipocyte activation through adrenergic-independent signaling.

### FOXP4 attenuates beige adipocyte thermogenesis by directly repressing *Pgc1α* and *Cebpβ* transcription

To explore the molecular mechanism underlying the impact of FOXP4 on beige adipocyte thermogenesis and activation, RNA-seq analysis were performed on beige adipocytes derived from sWAT-SVF of *Foxp4*_*AdipQ*_^Δ/Δ^ mice. As shown in Fig. 6A and B, expressions of an array of thermogenic or mitochondrial gene markers were elevated in *Foxp4*-deficient beige adipocytes. Next, we conducted ChIP-seq analysis of SVF-induced beige adipocytes by employing anti-FOXP4 and Anti-H3K27Ac antibodies for pulldowns. As shown in Fig. 6B, 5 common target genes were detected by both RNA-seq and ChIP-seq, including the classical thermogenic genes, *Pgc1α* and *Cebpβ*.

**Fig. 6.**
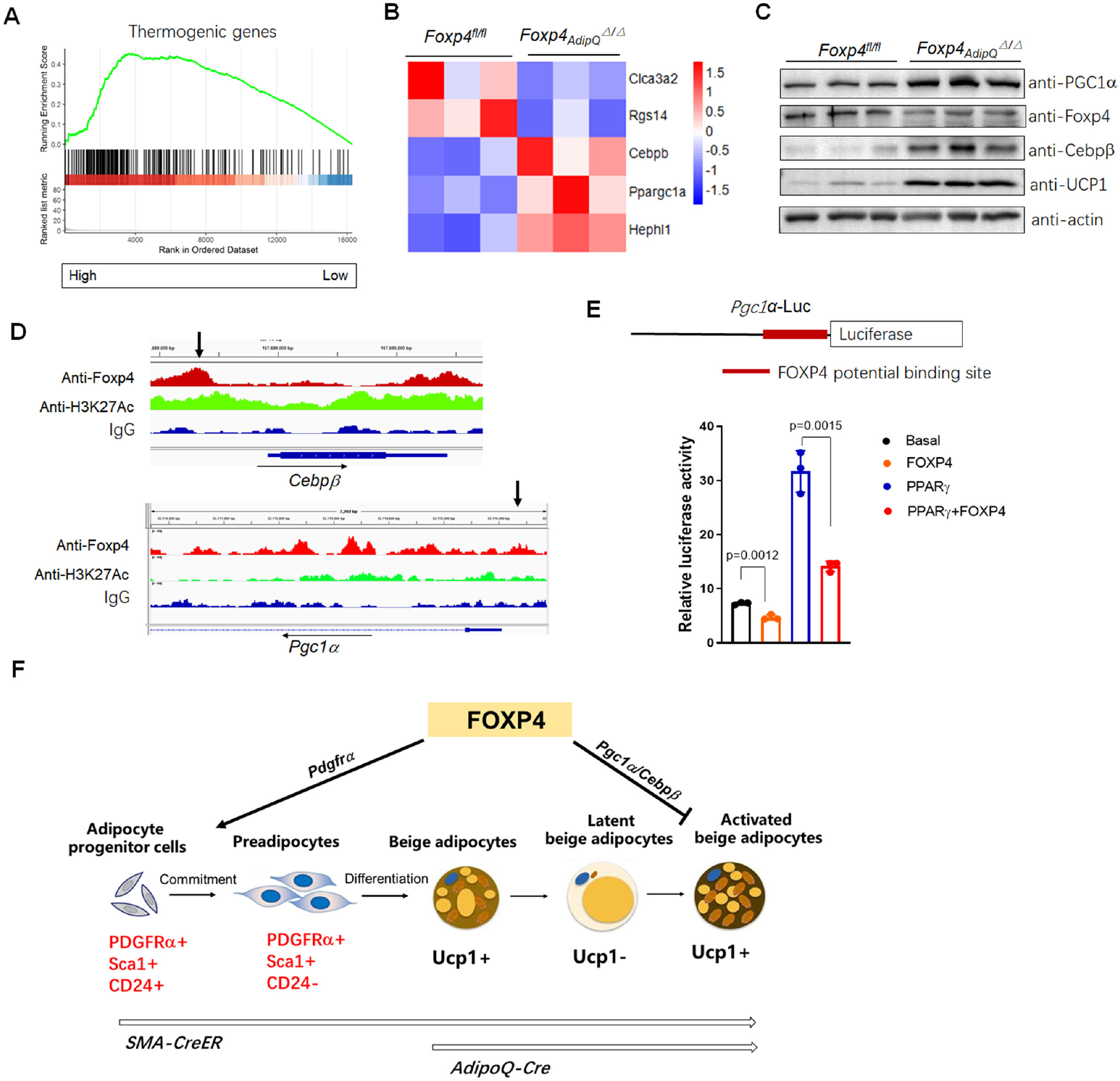
FOXP4 directly regulates the expressions of *Cebpβ* and *Pgc1α* in beige adipocyte thermogenic activation. A) RNA-seq analysis of thermogenic marker gene expressions in sWAT of *Foxp4*_*AdipQ*_^Δ/Δ^ mice under one-week 4 °C challenge. B) Heatmap of the 5 putative FOXP4-targeted gene expressions in (*A*). Chromatin occupancy analysis of ChIP-seq were conducted for SVF-derived beige adipocytes with anti-Foxp4 antibody. 5 common targets were detected to be overlapped with RNA-seq results. C) Western blot analysis showed the increase of protein expression of PGC1α, Cebpβ and UCP1 in sWAT of mice. D) ChIP-seq profile showed the FOXP4 binding sites (black arrows) within *Cebpβ* and *Pgc1α* promoter regions, which was consistent with anti-H3K27Ac binding domain. E) Luciferase reporter assay showed that FOXP4 repressed the transactivation of *Pgc1α*-Luc by PPARγ protein in 3T3-L1 cell lines. Upper panel depicts the construct of *Pgc1α*-Luc and potential FOXP4 binding site. F) Diagram depicting the distinct role FOXP4 in beige adipocyte differentiation and thermogenic activation. FOXP4 determines the beige/fibroblast cell fate choice in progenitor cells through modulating *Pdgfrα* signaling, whereas it suppresses their activation through repressing the expression of thermogenic genes *Pgc1α* and *Cebpβ*.

Western blotting confirmed the up-regulation of PGC1α, Cebpβ and UCP1 at the protein levels under conditions of decreased FOXP4 (Fig. 6C). Promoter occupancy as determined by ChIP-seq validated the relative enrichment of FOXP4 binding sites within the chromatin of *Pgc1α* and *Cebpβ* upstream promoter regions (arrows in Fig. 6D). In support, reporter assays employing a *Pgc1α* promoter-driven luciferase vector revealed that FOXP4 repressed the transactivation ability of PPARγ in 3T3-L1 cells (Fig. 6E). These findings demonstrated that FOXP4 restrained beige adipocyte thermogenic activation by directly repressing *Pgc1α* and *Cebpβ* gene transcription.

## DISCUSSION

Beige fat cells harbor distinctive molecular signatures from white and brown adipocytes during development. Once committed from progenitor cells, differentiated beige adipocytes demonstrate a plastic morphology that is tightly coupled with their reversible thermogenic potency. In this study, we dissected the phase-specific role of transcription factor FOXP4 in beige fat cell differentiation and thermogenesis. *Foxp4* deletion impaired the cell fate commitment of beige adipocytes from progenitor cells, but exacerbated cold-induced thermogenesis in differentiated beige adipocytes. Our findings indicate that FOXP4 primes beige cell differentiation, but acts as a “brake” for beige adipocyte thermogenesis under cold challenge (Fig. 6F).

To evaluate the impact of FOXP4 on beige cell differentiation, two independent knockout models were employed in our study. At the adult stage, the majority of cold-induced beige adipocytes arise from mural progenitor cells within vascular compartments of WAT (Long et al., 2014; Shamsi et al., 2021). It was previously shown that *SMA-Cre*^*ERT*^ mice could account for ∼60% of cold-induced beige adipocytes (Berry et al., 2016). In our first model, tamoxifen-inducible, Cre-mediated recombination was used to delete *Foxp4* in SMA^+^ SVF cells prior to beige adipocyte differentiation. Aside from SMA^+^ beige precursors, WAT-SVF cells contain another subset of PDGFRα-positive beige progenitor cells (Lee et al., 2012; Wang et al., 2013). As our second model we employed *pMSCV-Cre* to delete *Foxp4* in all beige precursor cells. Defective beige adipogenesis, as well as pro-fibrotic cell potency, was observed in progenitor cells with *Foxp4* deficiency. Of note, no defects of white adipocyte differentiation were observed at loss of *Foxp4* by *pMSCV-Cre* (Supplementary Fig. S2. E, F). These experiments provided compelling evidence demonstrating that FOXP4 was required for beige cell fate determination and differentiation.

PDGFRα-positive progenitor cells within sWAT harbor multiple potency in cell differentiation. They are precursor cells for both white and beige adipocytes (Gao et al., 2018), as well as for fibroblasts (Cattaneo et al., 2020). PDGFRα-positive progenitors are prone to give rise to beige adipocytes in response to β3-adrenoceptor activation and high-fat diet (Lee et al., 2012), rather than to cold induction (Berry et al., 2016; Shin et al., 2020). In addition, PDGFRα expression levels seem to be precisely controlled during adipogenesis. PDGFRα expression precedes beige adipocyte differentiation (Gao et al., 2018). Its continuous expression or activation in sequential phases drive progenitor cells toward fibroblastic cell fate (Iwayama et al., 2015; Marcelin et al., 2017; Sun et al., 2017). In line with those findings, *Foxp4*-deficient SVF cells display fibrotic signatures, due to increase and continuous *Pdgfrα* expression. When combined with our ChIP-seq and luciferase reporter data, the evidence suggests that FOXP4 primes a progenitor cell fate switch towards the beige adipocyte linage, partially through modulating *Pdgfrα* expression levels.

Prior to the present report, only a few beige-selective transcription factors have been characterized, including MRTFA (McDonald et al., 2015) and Tbx1 (Wu et al., 2012). FOXP4 is expressed in both BAT and sWAT of mice. But our genetic analysis showed that deletion of *Foxp4* in adipocytes had little effect on *in vivo* BAT development and thermogenesis. Only oxygen consumption and expression of several thermogenic marker genes were slightly exacerbated in *in vitro* cultures of SVF-derived brown adipocytes from *Foxp4*-deficient mice. Upon cold exposure and adrenoceptor agonist stimulation, we observed no significant differences *in vivo* in BAT thermogenesis. Its dynamic expression level in beige adipocytes during cell differentiation and cold exposure suggests that FOXP4 is a selective regulator of beige adipocyte development and cold-induced thermogenesis.

The thermogenic program of beige adipocytes can be activated through various pathways (Barbatelli et al., 2010; Rosenwald et al., 2013). Most beige adipocytes are activated through the β3-adrenergic signaling pathway (Lee et al., 2015). However, it also was reported that cold-induced activation of beige adipocytes requires the β1 adrenergic receptor (Adrb1), but not the β3 adrenergic receptor (Adrb3) (Jiang et al., 2017). Recently, a glycolytic beige population was identified that could be induced by chronic cold adaptation in the absence of β-adrenergic receptor signaling (Chen et al., 2019). Excess calorie intake also can trigger the activation of CHRNA2-dependent beige adipocytes (Jun et al., 2020), which also are glycolytic and β-adrenergic signaling independent. Interestingly, *Foxp4*-inactivated beige adipocytes appeared to only react to cold exposure, not to an adrenoceptor agonist (Supplementary Fig. S7C, D), and had elevated expressions of glycolytic marker genes as compared to controls in response to cold exposure (Supplementary Fig. S6C, D). This suggested that FOXP4 controls beige cell thermogenic activation through an adrenergic signaling-independent pathway, maybe partially through glycolytic pathway.

A recent report pointed out that different transcriptional machinery governs beige adipocyte development and activation between peri-weaning and adult stages (Wu et al., 2020). However, we observed that inactivation of *Foxp4* only slightly exacerbated UCP1-mediated thermogenesis at both periods. Thus, we suggest that FOXP4 is shared as a regulator of arresting thermogenesis in both juvenile and mature beige adipocytes. This view is consistent with previous studies from our laboratory that showed that FOXP1, a highly conserved paralogue of FOXP4, repressed both beige adipocyte differentiation and UCP1-mediated thermogenesis (Liu et al., 2019). Given that FOXP1 and FOXP4 form dimers in various tissues (Li et al., 2012; Li et al., 2004; Sin et al., 2015), we cannot exclude the potentially cooperative function of FOXP1/4 in controlling beige cell thermogenesis.

Collectively, the data presented in this report reveal a selective role of FOXP4 in beige adipocytes differentiation and cold-induced thermogenesis. We suggest that a more thorough understanding of the underlying causes of FOXP4 selective function will allow us to specifically manipulate beige cells to improve systemic energy metabolism and to combat obesity.

## MATERIAL AND METHODS

### Mice

The *Foxp4*^*fl/fl*^ constructed by our lab has been described in previous studies (Zhao et al., 2015). *Adiponectin-Cre* (Stock no. 028020 in Jax Lab) was obtained from Jax lab. *SMA-CreERT* mice was kindly provided by Prof. Gang Ma in Shanghai Jiaotong University. The genetic backgrounds of all knockout mice were C57Bl/6J. Mice were bred with standard rodent chow food or HFD. For cold treatment, mice were bred under 4 °C environment for a week. For continuous β-adrenergic stimulation, mice were intraperitoneal injected CL-316,243(0.75 mg/kg) for up to 7 days. Male mice were used in the experiments unless otherwise indicated. The experiments were not randomized, and the investigators were not blinded to allocation during experiments or outcome assessments. Mice are maintained under a constant environmental temperature (22 °C) and a 12-h light/12-h dark cycle. All animal experiments were performed according to the guidelines of Shanghai Jiao Tong University (SYXK 2011-0112).

### Metabolic study

Minispec TD-NMR Analysers (Bruker Instruments) were used to evaluate adiposity composition on anesthetized animals. Food intake, energy expenditure, O2 consumption, CO2 production and physical activity were measured by using indirect calorimetry system (Oxymax, Columbus Instruments), installed under a constant environmental temperature (22 °C) and a 12-h light (07:00–19:00 hours), 12-h dark cycle (19:00–07:00 hours). Mice in each chamber had free access to food and water. The raw data were normalized by body weight and the histograms of day (07:00-19:00 hours) and night (19:00 – 07:00 hours) values were the mean value of all points measured during the 12-h period.

### Immunohistochemistry and TEM

Adipose tissues were fixed in 4% PFA for 16 hours at 4 °C, embedded in paraffin or tissue freezing medium (Leica) and sectioned to 4 μm. H&E staining was conducted according to standard protocols. For immunofluorescence, heat-induced antigen retrieval with sodium citrate buffer (10mM sodium citrate, 0.05% Tween 20, pH 6.0) was performed before sections were blocked with 5% BSA in TBST (pH 7.6) for 30 minutes at 37 °C, then incubated overnight at 4 °C with primary antibodies to mouse Foxp4 (Millipore, #ABE74, 1:100), UCP1 (Abcam, #ab10893, 1:100). Subsequently, sections were incubated with secondary fluorescent-conjugated or HRP-conjugated antibodies at 37 °C for 30 minutes in the dark. Samples were imaged by the Leica TCS SP5 confocal microscope, Leica DM2500, or Leica 3000B microscope. Transmission electron microscopy (TEM) of beige adipose tissue was carried out in accordance with a previous study (Liu et al., 2019).

### Cell cultures

For SVF cell isolation, primary BAT and sWAT were digested with 1 mg ml^−1^ collagenase type I (Sigma) in DMEM (Invitrogen) supplemented with 1% bovine serum albumin for 25 min at 37 °C, followed by filtration, density separation with centrifugation. The freshly isolated SVF cells were seeded and cultured in growth medium containing DMEM, 20% FBS, 1% penicillin/streptomycin (P/S) at 37 °C with 5% CO_2_ for 3 days, followed by feeding with fresh medium every 2 days to reach confluence. For brown/beige adipocyte differentiation, the cells were induced with induction medium contains DMEM, 10% FBS, 5 μg ml^−1^ insulin, 0.5 mM isobutylmethylxanthine (Sigma), 1 μM dexamethasone (Sigma), 50 nM T3 (Sigma) and 5 μM troglitazone (Sigma) for 48 hours, and further in growth medium supplemented with insulin, T3 and troglitazone for 6 days, followed by 0.5 mM cyclic AMP (Sigma) treatment for another 4 hours. For inducible knockout in beige cells, SVF from *Foxp4*_*SMA*_^*ErtΔ/Δ*^ and *Foxp4*^*fl/fl*^ mice were expanded for 2 days with growth medium, treated with tamoxifen in cultures for 24 hours with final concentration 1nM. followed by 6-day differentiation cultures without tamoxifen. HEK293T (ATCC) were cultured in DMEM with 10% FBS. For oil red staining, cultured cells were washed with PBS and fixed with 10% formaldehyde for 15 min at room temperature. Then the cells were stained using the Oil red O working solutions (5g/l in isopropanol) and 4 ml H_2_O for 30 min. After staining, the cells were washed with 60% isopropanol and pictured.

### FACS analysis

2-month-old *Foxp4* knockout mice by *SMA-CreERT* were interperitoneally injected once with tamoxifen dissolved in sunflower oil (Sigma, 100 mg/kg). After 48 hours, SVF cells were isolated from mice euthanized with CO_2_ according to protocols showed above. Progenitor cells from SVF were expanded for two days in growth cultures and treated with tamoxifen (a final concentration of 1 nM) for 24 hours. Then those cells were treated with fresh cultures without tamoxifen for another 24 hours before being collected for FACS analysis. Tamoxifen-administrated SVF cells were suspended with FACS buffer to a final concentration of 10^5^-10^6^ cells/100uL, incubated on ice for 30∼45 minutes with antibody combinations of PerCP-CD45 (BioLegend, #103131), PerCP-CD31 (BioLegend, #102419), APC-Sca-1 (BioLegend, #122511), PE-CD140α (BioLegend, #135905), CD24-FIFC (BioLegend, #101815). The cells then washed twice with PBS before analysis on BD Calibur. Proportion of adipocyte progenitors were analyzed with Flowjo V10 software.

### Oxygen consumption assay

Primary SVF cells from BAT and sWAT were isolated and cultured for 4 days before being plated in XF cell culture microplates (Seahorse Bioscience). SVF cells (10,000 cells) were seeded in each well, and each treatment included cells from three BAT or sWAT replicates. After 6-day differentiation, cultured adipocytes were washed twice and pre-incubated in XF medium (supplemented with 25mM glucose, 2mM glutamine and 1mM pyruvate) for 1–2 h at 37 °C without CO_2_. The OCR was measured using the XF Extracellular Flux Analyser (Seahorse Biosciences). Oligomycin (2 mM), FCCP (2 mM), and Antimycin A (0.5 mM) were preloaded into cartridges and injected into XF wells in succession. OCR was calculated as a function of time (pmoles per minute per μg protein).

### Luciferase assay

Luciferase assays were performed in HEK293T or 3T3-L1 cells. The reporter plasmid, *Pgc1α-Luc* containing a 2.6 kb fragment of the promotor region of the *Pgc1α* gene, was obtained from Dr. JiQiu Wang of the Ruijin hopital (Shanghai, China). *Pdgfrα-Luc* plasmid containing 1.3 kb of the 5’ flanking region of *PDGFRα* gene was constructed by our lab. The primers used for amplification are shown in Table S1. The expression plasmids of *Foxp4* and *Pparγ* were constructed into the pcDNA3.0 vector. Cells were transfected using FuGENE HD (Promega) in 24-well plates. The transfection amount of each plasmid was 200 ng, and the total amount of transfected DNA across each transfection was balanced by pcDNA3.0 plasmids when necessary. After 36 hours, dual luciferase assay was performed according to the manufacturer’s protocols (Promega).

### RNA isolation and quantitative RT-PCR

We used TRIzol (Vazyme, #R401) and for total RNA extraction, respectively, according to the manufacturers’ instructions. Extracted RNA (1μg) was converted into cDNA using the HiScript® III SuperMix for qPCR (Vazyme,R323-01). Quantitative RT-PCR (qRT-PCR) was performed using an LightCycler® 480 II (Roche) and SYBR Green PCR Master Mix (Vazyme, #Q711-02). Fold change was determined by comparing target gene expression with the reference gene β-actin. The primers used for qRT-PCR are shown in Table S1. For RNA-seq, total RNA extracted from sWAT with TRIzol was used for library construction and RNA sequencing (Personal Biotechnology Co., Ltd, Shanghai, China).

### Western blot

For western blotting, adipose tissue was homogenated and lysed with RIPA (Beyotime, P0013B). Protein samples were incubated with primary antibodies against Foxp4 (Millipore, #ABE74, 1:1000), Ucp1 (Abcam, #ab10893, 1:1000), C/ebpβ (Santa Cruz, #sc-150, 1:500), PGC1α (Abways, #CY6630,1:1000), β-actin (Selleck, #A1016, 1:2000) at 4 °C overnight. Proteins were visualized using HRP-conjugated secondary antibody and chemiluminescent HRP substrate (Millipore).

### ChIP-Seq

Wash 20μl Magna ChIP™ Protein A+G Magnetic Beads (Millipore, 16-663) twice with 1 ml FA buffer (10mM HEPES[PH7.5],150mM NaCl, 1mM EDTA, 1% TritonX-100, 0.1%Sodium deoxycholate, 0.1% SDS and protease inhibitors). Then suspend the beads with 1 ml FA buffer, add 4ug antibody to the beads and rotate for at least 2 hours. Differentiated SVF cells were cross-linked using 1% formaldehyde in PBS at room temperature with rotation. Cells were then incubated with 125mM Glycine (Sangon, #A610235-0500)(62.5ul 2M Glycine/ml PBS)at room temperature for 10 minutes to stop cross-linking. After washed twice with ice-cold PBS, cells were collected and diluted in 0.5 ml FA buffer. Sonication the cells by sonics CV130 with the parameter: 5s on;10s off; 6 minutes to make the DNA fragment at 300-500bp. Centrifuge the sonicated solution at 13,000 rpm for 5 minutes to Collect the supernatant. Wash the beads bounded with antibody twice with FA buffer. Add antibody coated beads into the sonicated supernatant for 12-16 hours at 4 °C with rotation. Wash the beads sequential with FA buffer once, high salt buffer (10mM HEPES[PH7.5], 150mM NaCl, 1mM EDTA, 1% TritonX-100, 0.1%Sodiumdeoxycholate, 0.1%SDS), LiCl buffer (10mM Tris-HCl[PH8.0], 0.25M LiCl, 1% NP-40, 1mM EDTA, 0.1%Sodiumdeoxycholate) and TE buffer (10mM Tris-HCl[PH7.5], 1mM EDTA) twice. Suspend the beads with 270ul Elution buffer (50mM Tris-HCl[PH7.5], 1mM EDTA, 1% SDS) and elute it with 68 °C, 900 rpm for 30minutes on an eppendorf thermoMixer. After elution, add 130 µl TE buffer, 3 µl Rnase A (Thermo fisher, EN0531) into the eluate, incubate at 37 °C for 30minutes. Then add 5ul Proteinase K (Thermo fisher, #AM2546),incubate at 65 °C overnight to reverse cross-links. DNA was isolated and purified with a ZYMO DNA clean & Concentrator (ZYMO RESEARCH, #D4013).

Libraries were constructed using an VAHTS Universal DNA Library Prep Kit for Illumina® V3(Vazyme, #ND607) according to the manual protocol. Isolated Chip DNA was sequential subjected to end repair/phosphorylation/A-tailing adding, and index adaptor ligation. AMPure XP beads (Beckman Coulter, #A63880) were used for a post-ligation cleanup, DNA was eluted from beads and amplified by PCR for 10 cycles. DNA sized between 200 and 700bp was selected by a double-sided size selection strategy with AMPure XP beads. After elution with 10mM Tris[PH8.0],libraries were analyzed using the Qubit and Agilent 2100 Bioanalyzer, pooled at a final concentration of 12pM and sequenced on a HiSeq2500. For ChIP-seq analysis, demultiplexed ChIP-seq reads were aligned to the mm10 mouse genome using Bowtie2 (45) with the parameter “--no-discordant --no-unal --no-mixed”. PCR duplicates and low-quality reads were removed by Picard. Reads were processed using Samtools (46)and subjected to peak-calling with MACS2 (47) with a parameter “except -f BAMPE -p 0.01”. We convert the bam file to bw file using deeptools (v3.3.0). Integrative Genomics Viewer (IGV, v2.7.2) was used for peak visualization. Overlaps between Chip-seq and RNA-seq were performed and we draw the venn diagram in R.

### Glucose tolerance test (GTT)

For GTT, mice were given i.p. injection of 100 mg/ml D-glucose (2 g/kg body weight) after overnight fasting, and tail blood glucose concentrations were measured by a glucometer (AccuCheck Active, Roche).

### Data analysis

For RNA-seq analysis, sequencing reads were filtered using Cutadapt and aligned to the mm10 mouse genome using HISAT2 (Kim et al., 2015). Filtered reads were assigned to the annotated transcriptome and quantified using HTSeq (Anders et al., 2015), FPKM was used as normalization method. The analysis below was all performed in R. We used DEseq Package for differential expression analysis (Anders and Huber, 2010). Genes were considered significant if they passed a fold change (FC) cutoff of |log2FC|>1 and a false discovery rate (FDR) cutoff of FDR%<0.05. Heatmap package was used for gene expression cluster analysis and heatmap visualization. We used topGO for GO analysis (Alexa et al., 2006). Clusterprofiler package was used for KEGG and GSEA analysis with default parameter (Yu et al., 2012).

For statistical analysis, all data are presented as mean ± SD. Error bars are SD. Two-tailed Student’s t-tests for comparisons between two groups and two-way ANOVA for that more than two groups.

## ACKKNOWLEDGEMENT

This work was supported by research grants from the National Natural Science Foundation of China [92068203, 91749103, 81421061, 31100624 and 81200586] and grant from the National Major Fundamental Research 973 Program of China [2020YFA0803601, 2014CB942902] to X.G. Great thanks for Prof. Haley O. Tucker from University of Texas in manuscript editing.

## AUTHOR CONTRIBUTIONS

F.W., S.X., T. C., W.Z., S.L, S. W., R. Z., X.X. performed experiments, X.G. designed experiments, P. L., X.Z., Z.Y. and J.W. helped preparing samples and instructing experiments, X.G. wrote manuscript.

## DATA AND RESOURCE AVAILABILITY

The data sets generated during and/or analyzed during the current study are available from the corresponding author upon reasonable request. The resources generated during and/or analyzed during the current study are available from the corresponding author upon reasonable request.

## Supplementary figures and figure legends

**Fig. S1.**
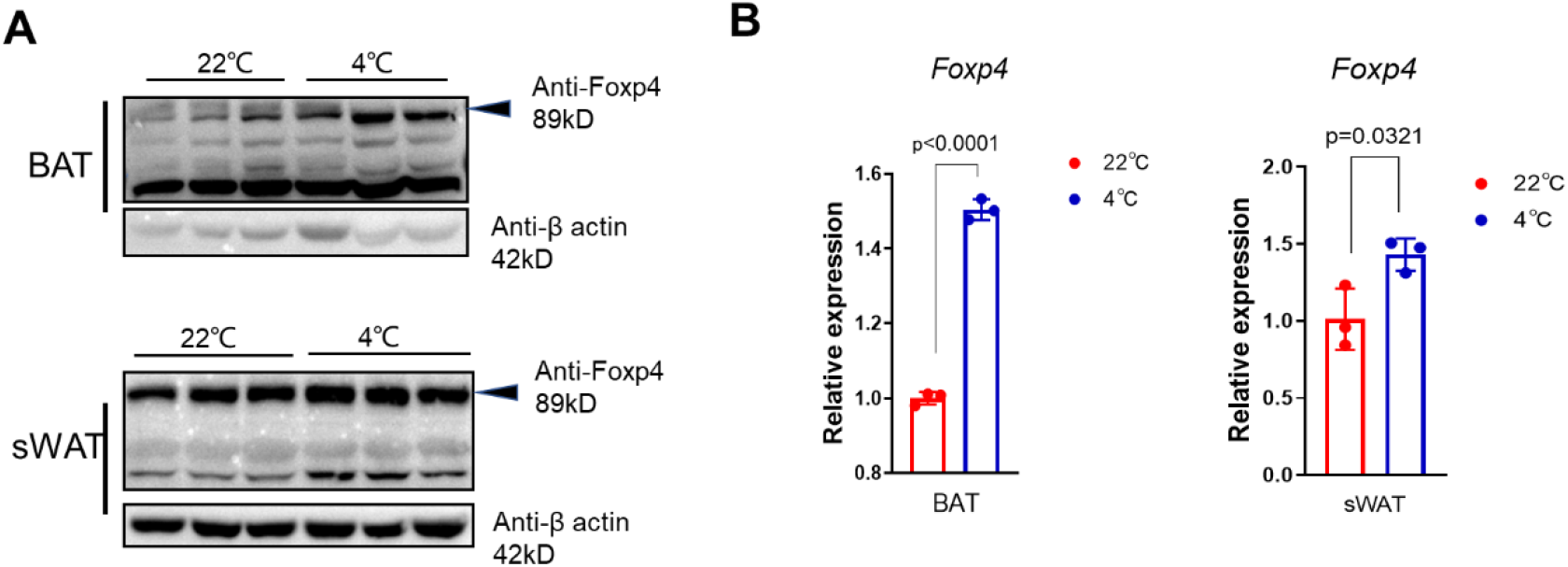
Expression of *Foxp4* in adipose tissues from *Foxp4*_*AdipQ*_^Δ/Δ^ mice. A) Western blot for FOXP4 protein in BAT and sWAT of mice under room temperature (22 °C) and one-week cold exposure (4 °C). B) mRNA levels of *Foxp4* expression in BAT and sWAT from mice of (A). n, 3.

**Fig. S2.**
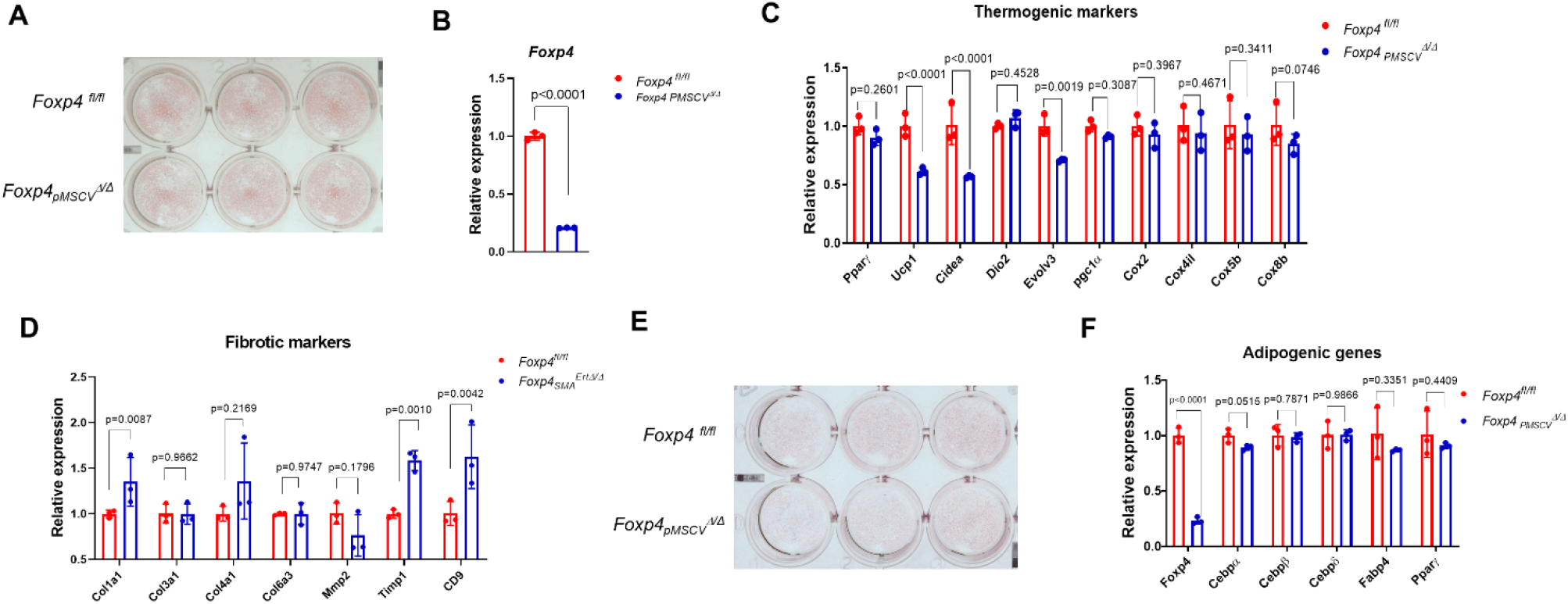
*In vitro* brown adipocyte differentiation with *Foxp4* deficiency by *pMSCV-Cre*. A) Oil Red O staining for 8-day brown adipocyte differentiation from *pMSCV-Cre*-transfected SVF of BAT from *Foxp4*^*fl/fl*^ mice. B, C) mRNA levels of *Foxp4*, thermogenic and brown selective markers in cells of (A). n, 3. C) mRNA levels of fibrotic cell marker genes in beige differentiation from SVF of *Foxp4*_*Sma*_^*ErtΔ/Δ*^ mice with *Foxp4* inactivation induced by tamoxifen in cultures. D) Oil Red O staining for 8-day white adipocyte differentiation from *pMSCV-Cre*-transfected SVF of sWAT from *Foxp4*^*fl/fl*^ mice. E) mRNA levels of adipogeniesis markers in cells of (A). n, 3.

**Fig. S3.**
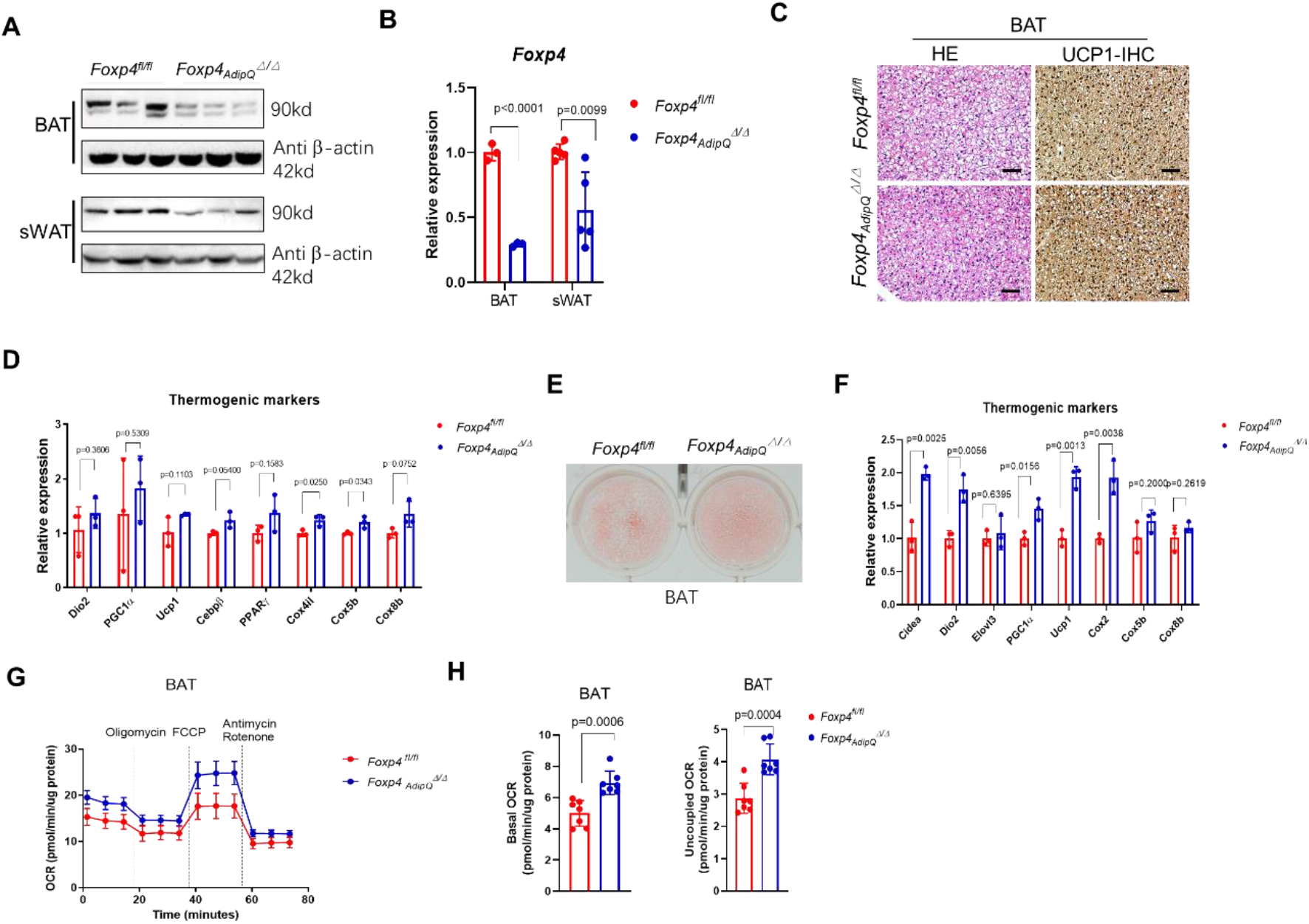
Thermogenesis in BAT of *Foxp4*_*AdipQ*_^Δ/Δ^ mice. A) Western blot for FOXP4 protein in BAT and sWAT of *Foxp4*^*fl/fl*^ and *Foxp4*_*AdipQ*_^Δ/Δ^ mice at age of 2 months old. (B) Assessment of *Foxp4* mRNA expression in BAT and sWAT from mice by qPCR. n, 3. C) H&E and immunohistochemical staining (IHC) for UCP1 on BAT sections from *Foxp4*_*AdipQ*_^Δ/Δ^ mice. D) mRNA levels of thermogenic and mitochondrial markers in BAT. E) Oil Red O staining 8 day post brown adipocyte differentiation from BAT-SVF of *Foxp4*^*fl/fl*^ and *Foxp4*_*AdipQ*_^Δ/Δ^ mice at age of 8 weeks. F) mRNA levels of thermogenic markers for brown adipocytes in (E). n, 3. G) Oxygen consumption rate (OCR) was measured for brown adipocytes from (E). Uncoupled respiration was recorded after oligomycin inhibition of ATP synthesis, and maximal respiration following stimulation with carbonyl cyanide 4-(trifluoromethoxy) phenylhydrazone (FCCP). n, 3. H) Quantitative analysis of basal and uncoupled OCR in (G). n, 3.

**Fig. S4.**
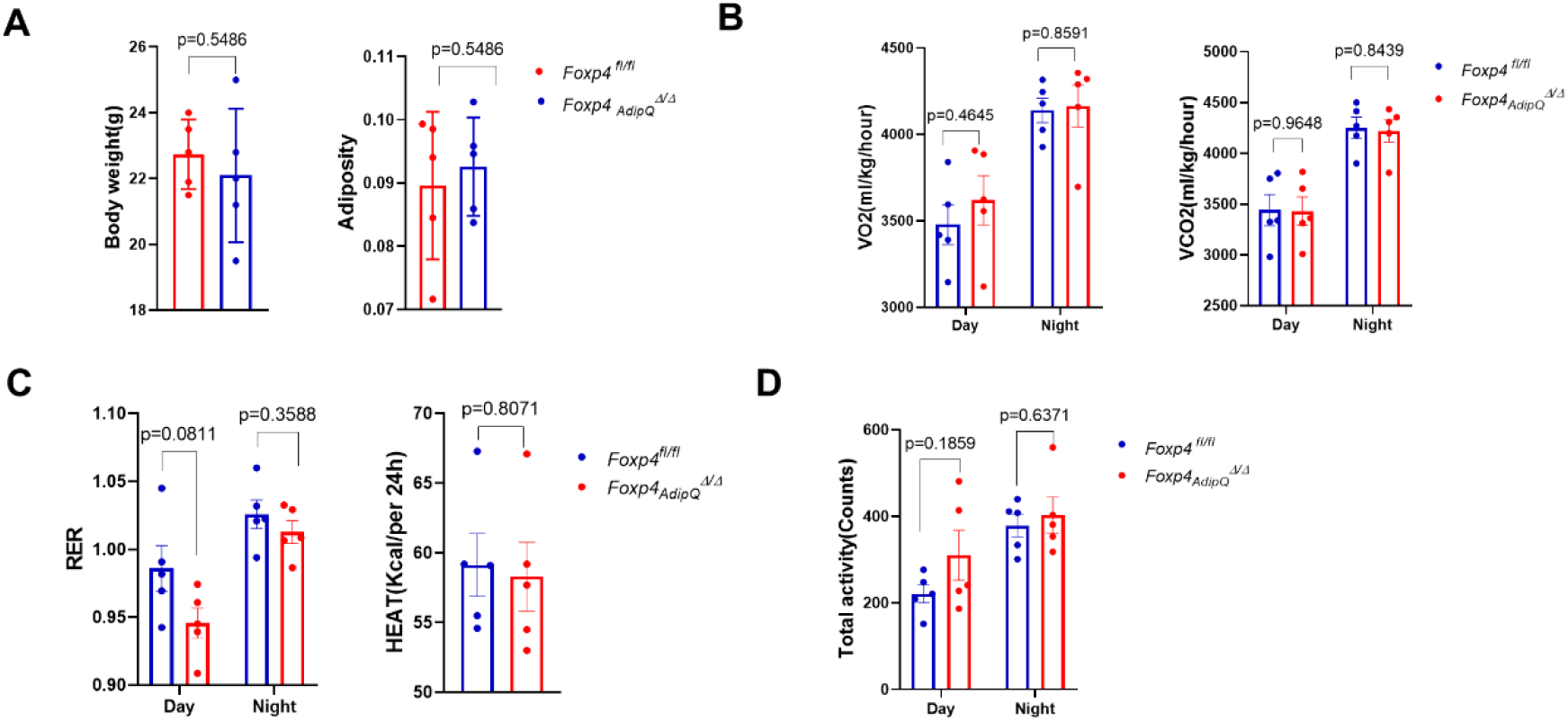
Metabolic analysis for *Foxp4*_*AdipQ*_^Δ/Δ^ mice. A) Body weight and relative adiposity of *Foxp4*^*fl/fl*^ and *Foxp4*_*AdipQ*_^Δ/Δ^ mice during day and night in metabolic cages at age of 3 months old. n, 5. B) Quantification of O_2_ and CO_2_ consumption of mice under room temperature. n, 5. C) RER and heat production of mice. n, 5. D) Total activity of mice. n, 5.

**Fig. S5.**
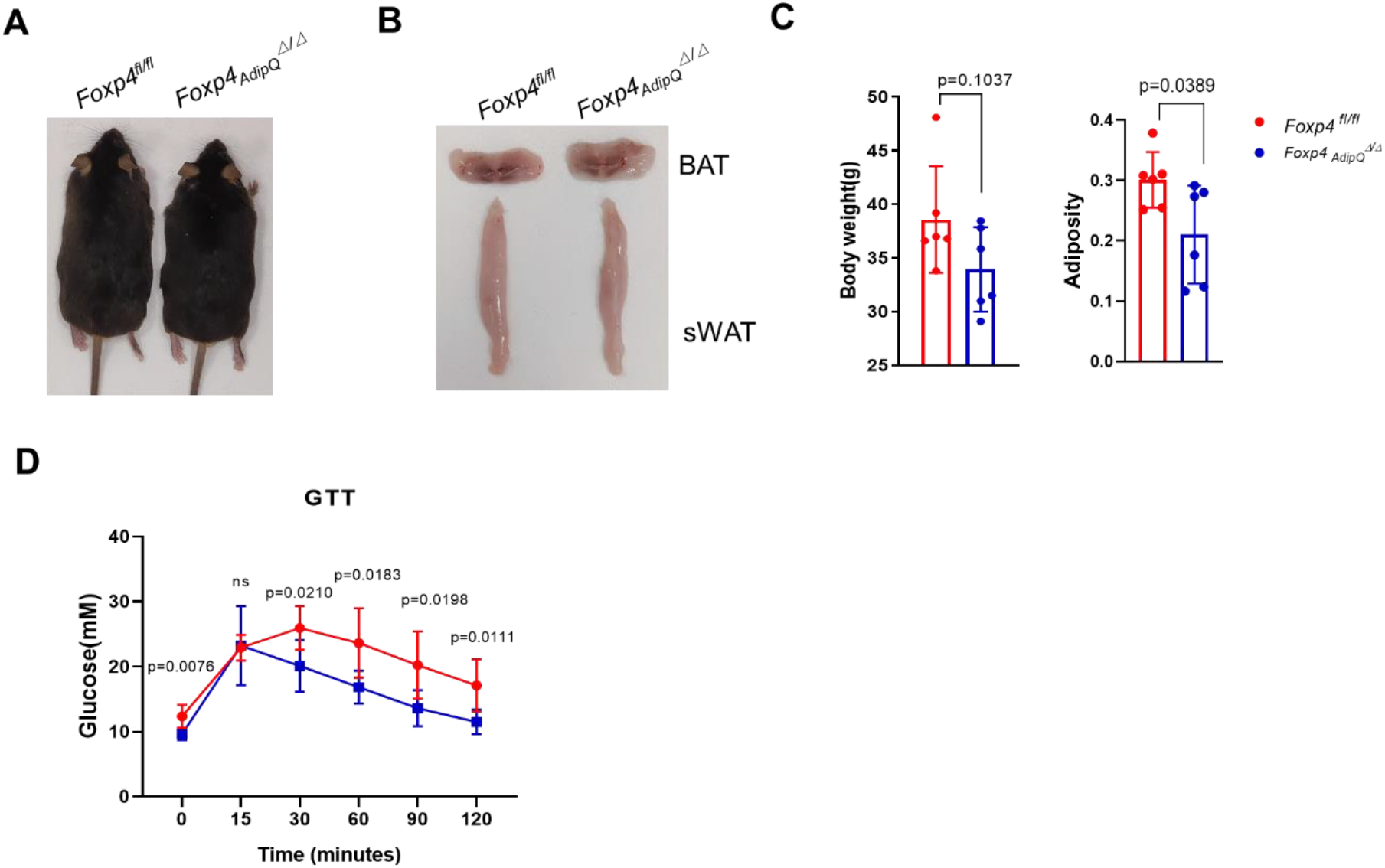
*Foxp4* deficiency protects mice from HFD-fed obesity. A) Dorsal view of representative *Foxp4*^*fl/fl*^ and *Foxp4*_*AdipQ*_^Δ/Δ^ mice of after 8-week feeding with HFD at age of 2 months. B) Representative fat depot of BAT and sWAT from mice of (A). C) Body weight and relative adiposity of HFD-fed mice of (A). n, 6. D) GTT of HFD-fed mice. n, 6.

**Fig. S6.**
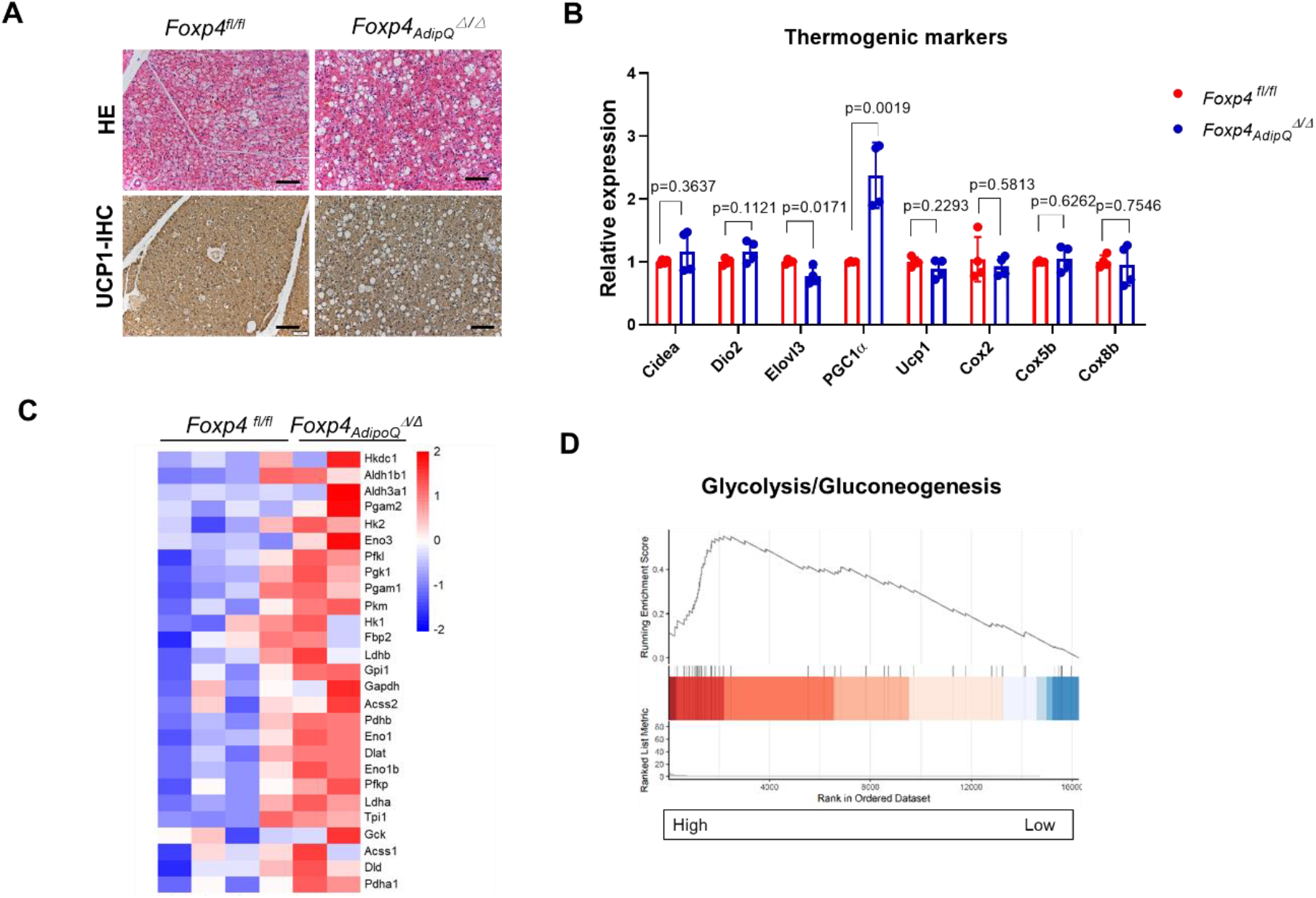
BAT thermogenesis in *Foxp4*_*AdipQ*_^Δ/Δ^ mice upon cold exposure. A) H&E and immunohistochemistry (IHC) staining for UCP1 protein on BAT sections from *Foxp4*_*AdipQ*_^Δ/Δ^ mice after one-week cold exposure at 4 °C. B) Thermogenesis in BAT of (A) assessed by qPCR with selective markers (*Cidea, Dio2, Elovl3, PGC1α, Ucp1, Cox2, Cox5b, Cox8b*). n, 3. C) Heatmap depicting the mRNA levels of glycolytic genes in beige adipocytes from sWAT in *Foxp4*_*AdipQ*_^Δ/Δ^ mice after one-week cold exposure at 4 °C. D) RNA-seq analysis of glycolytic marker gene expressions in sWAT of (C).

**Fig. S7.**
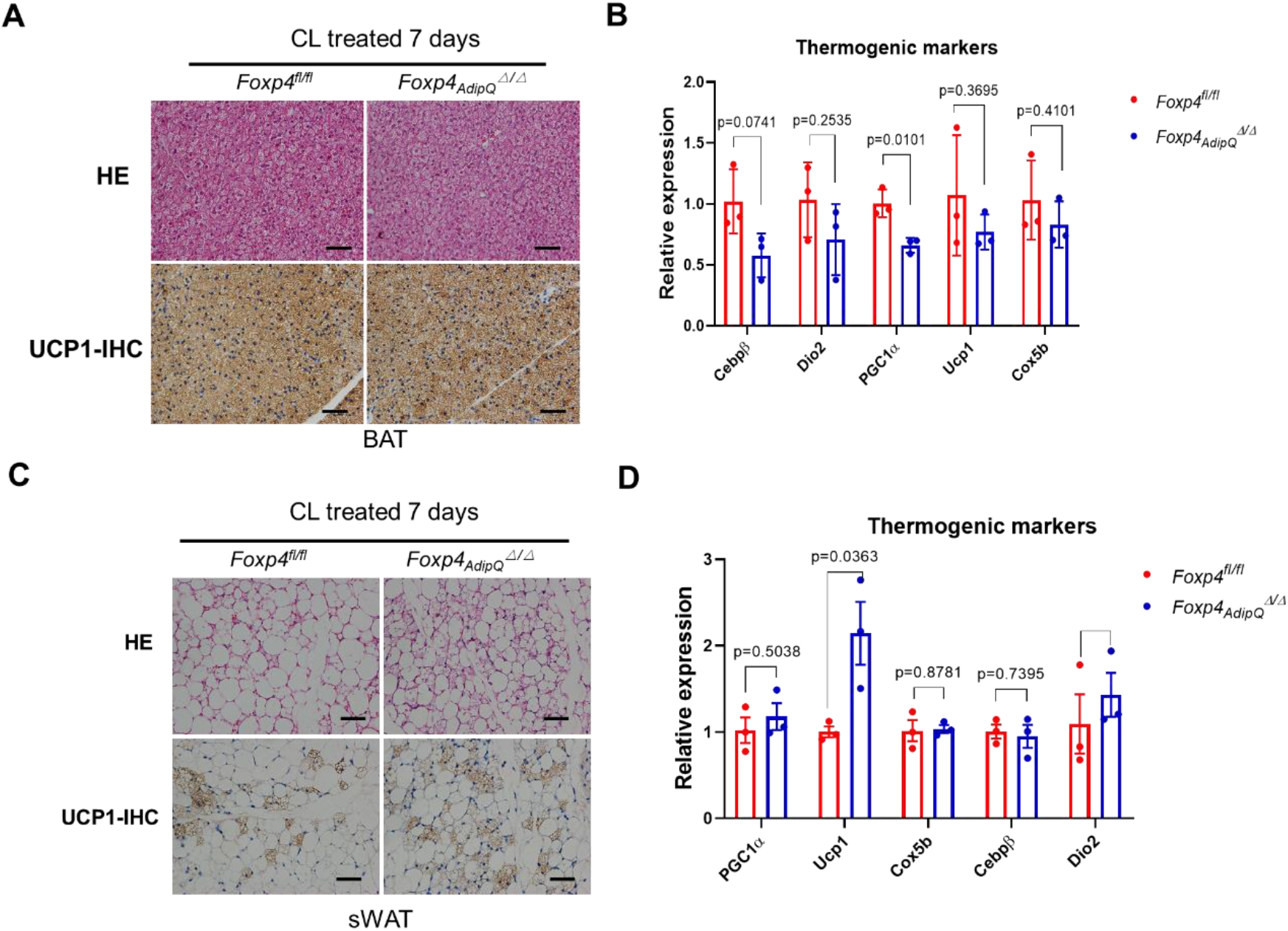
Thermogenic activation in CL-316,243-stimulated *Foxp4*_*AdipQ*_^Δ/Δ^ mice. A, C) H&E and IHC staining for UCP1 on BAT (A) and sWAT (C) sections from mice stimulated with CL-316,243 for 7 days. B, D) qPCR analysis for thermogenic gene expressions in BAT (B) and sWAT (D) from CL-316,243-stimulated mice. n, 5.

**Supplementary table S1.**
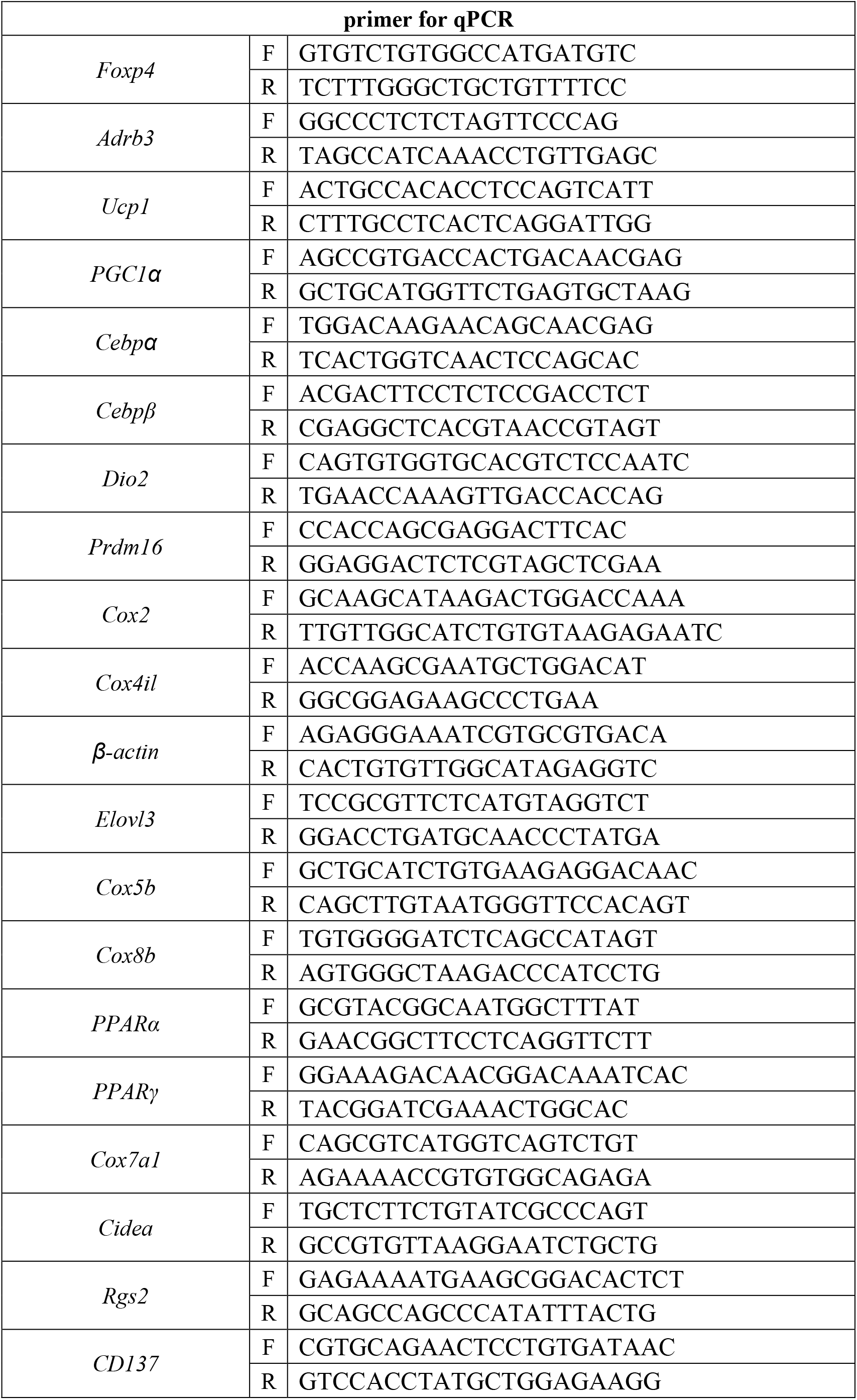
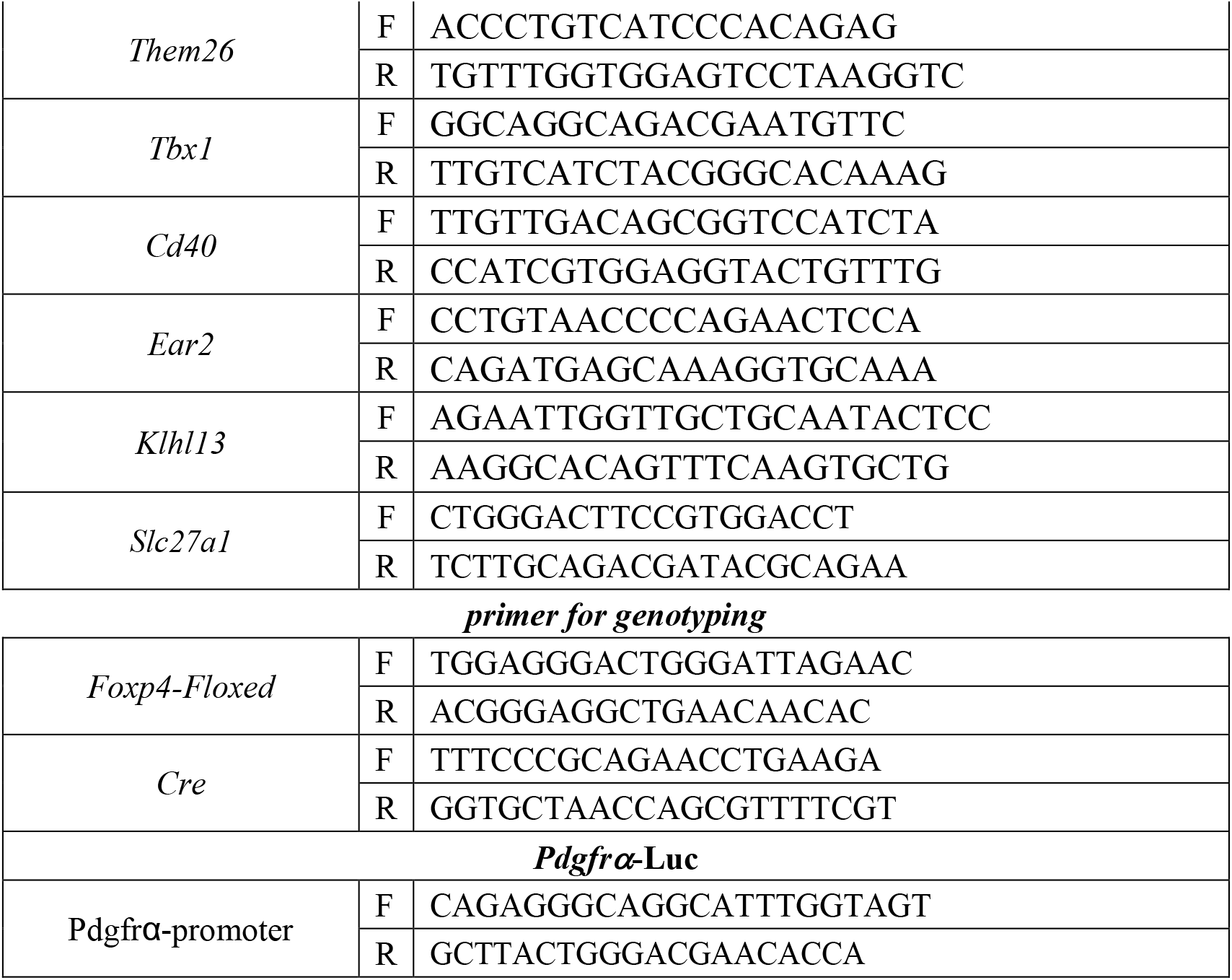
primers for qPCR and genotyping.

